# Genetic Interaction between RLM1 and F-box Motif Encoding gene SAF1 Contributes to Stress Response in *Saccharomyces cerevisiae*

**DOI:** 10.1101/757773

**Authors:** Meenu Sharma, V. Verma, Narendra K Bairwa

## Abstract

Stress response is mediated by transcription of stress responsive genes. F-box motif protein Saf1 involves in SCF-E3 ligase mediated degradation of the adenine deaminase, Aah1 upon nutrient stress. Four transcription regulators, BUR6, MED6, SPT10, SUA7, have been reported for SAF1 gene in genome database of *Saccharomyces cerevisiae.* Here in this study an *in-silco* analysis of gene expression and transcription factor databases was carried out to understand the regulation of SAF1 gene expression during stress for hypothesis generation and experimental analysis. The GEO profile database analysis showed increased expression of SAF1 gene when treated with clioquinol, pterostilbene, gentamicin, hypoxia, genotoxic, desiccation, and heat stress, in WT cells. SAF1 gene expression in stress conditions correlated positively whereas AAH1 expression negatively with RLM1 transcription factor, which was not reported earlier. Based on analysis of expression profile and regulatory association of SAF1 and RLM1, we hypothesized that inactivation of both the genes may contribute to stress tolerance. The experimental analysis with the double mutant, *saf1Δrlm1Δ* for cellular growth response to stress causing agents, showed tolerance to calcofluor white, SDS, and hydrogen peroxide. On the contrary, *saf1Δrlm1Δ* showed sensitivity to MMS, HU, DMSO, Nocodazole, Benomyl stress. Based on in-*silico* and experimental data we suggest that SAF1 and RLM1 both interact genetically in differential response to genotoxic and general stressors.

## Introduction

The natural population of eukaryotic cells, mostly microorganism remains in non-dividing state and proliferate upon nutrients availability (GRAY *et al.* 2004).The proliferation of *Saccharomyces cerevisiae* cells from in and out of quiescence phase due to nutrients availability or stress condition remains an active research area for both basic research and biotechnological purpose. Upon nutrient deprivation yeast cells stop dividing and enter into a stationary phase. This transition leads to glycogen accumulation and reduced cell wall porosity which makes cells to resist stressor which increases survival probability. Mutant cell which are not able to enter into stationary phase showed sensitivity to stress condition and may die due to starvation (WERNER-WASHBURNE *et al.* 1993).Gene expression studies with WT cells on entry into stationary (DERISI *et al.* 1997) phase, showed varied expression of thousands of genes (LASHKARI *et al.* 1997; GASCH *et al.* 2000). These studies implicate that active transcriptional regulation is necessary for entry or exit from stationary phase. The phase transition process is dependent on the ubiquitin proteasome system as it was reported that ubiquitin is required for survival during starvation (FINLEY *et al.* 1987). *S. cerevisiae*, SCF-E3 ligase component Saf1, recruits Aah1, adenine deaminase for degradation upon nutrient deprivation condition. Mechanism of Aah1 degradation associated with entry or exit from quiescence stage is interesting gene model to understand the stress and quiescence phase. The nutrient deprivation condition induces stress which lead cell to enter into the quiescence phase. The transition from actively dividing cells to quiescence phase leads to changes in the gene expression and transcription regulation. AAH1 gene expression is down regulated during transition from proliferative state to quiescence state by Srb10p and Srb11p at transcriptional level. AAH1 encodes adenine deaminase which converts adenine to hypoxanthine. At post-transcriptional level Aah1p is degraded by Saf1 mediated proteasomal process. The SCF-Saf1 pathway has been reported in the degradation of the unprocessed vacuole and lysosomal proteins (MARK *et al.* 2015). The serine proteases of the vacuole play a major role during starvation of a cell. In vacuolar region three proteins were reported named as serine protease B (PRB1), protease C (PRC1), and Ybr139w target for proteasomal degradation by Saf1 (MARK *et al.* 2015). A comprehensive study on binding of gene regulatory protein by chromatin immunoprecipitation-chip analysis upon heat treatment showed three regulatory transcription factors (BUR6, MED6, SUA7) for the SAF1 gene (VENTERS *et al.* 2011). The Bur6 (YER159C) forms a complex with NC2 transcription regulator, which is heterodimeric in nature. The Med 6 (YHR058C) forms the part of RNA polymerase II mediator complex which is essential for transcriptional regulation. The expression of MED6 elevated during DNA replication stress. The Sua7 (YPR086W), a general transcription factor, is required for transcription initiation by RNA polymerase II. In a microarray RNA expression based study, SPT10 histone H3 acetylase `acts as` transcriptional regulator of the SAF1 gene (MENDIRATTA *et al.* 2006). The null mutant of SAF1 showed the decreased death rate at elevated temperature (TENG *et al.* 2011) and resistance to histone deacetylase inhibitor CG-1521 (GAUPEL *et al.* 2014).

RLM1/YPL089C (resistance to lethality of MKK1P386 over expression) is a MADS-box (Mcm1, Agamous, Deficiens, and serum response factor) transcription factor is phosphorylated and activated by the MAP-kinase Slt2p (WATANABE *et al.* 1997). The *Saccharomyces cerevisiae* cells treated with calcofluor white and zymolyase showed the activation of Slt2p MAP kinase pathway and Rlm1 as final signalling molecule regulated the expression of genes related to cell wall integrity (BOORSMA *et al.* 2004).

Gene expression omnibus database (GEO) at the National Center for Biotechnology Information (NCBI) is repository of genome wide gene expression data submitted by different laboratories on different model organism. The database allows the mining of microarray based gene expression information on model organism using both experiment-centric and gene-centric approach (BARRETT and EDGAR 2006). The GEO profile can be checked as gene name or ID across all the experimental condition and experiment type (BARRETT *et al.* 2005). Gene expression is dependent on the transcriptional network which relates to the coordinated expression, involving interaction of transcription factors and promoters of multiple genes (HUGHES and DE BOER 2013). The Yeast transcription database YEASTARCT (Yeast Search for Transcriptional Regulators And Consensus Tracking) allows the search of association between the transcription factors and gene expression in variety of experimental condition (http://www.yeastract.com). It is repository of curated association between transcription factors and genes in *Saccharomyces cerevisiae* (TEIXEIRA et al. 2018). DNA microarray technology has been useful which measure the gene expression in response to stress caused by environmental changes (WANICHTHANARAK *et al.* 2014). It also allows understanding of transcriptional regulations during stress. The Yeast stress expression database (yStreX: http://www.ystrexdb.com/) is source for analysis of gene specific expression pattern during stress. The database has collection of genome wide expression data on stress response in *S. cerevisiae* (WANICHTHANARAK *et al.* 2014).

Here we have studied the Gene Expression Omnibus profile database (GEO) for expression status of SAF1 gene during stress conditions in WT cells. Yeast transcription databases were investigated for association of transcription factors with the SAF1 expression during stress. The regulatory association analysis was done to understand, transcriptional regulation of target genes by transcription factors during stress. Further, yeast stress expression database (yStreX) was also analysed the expression status of SAF1 and their associated transcription factor upon stress to generate hypothesis and validated by experimental. Here, we report RLM1 as novel transcription factor regulating expression of SAF1 during stress through *in-silico* analysis. Further, we hypothesized that ablation of both the genes may contribute to stress tolerance. We report the loss of SAF1 and RLM1 together leads to stress resistance in *S. cerevisiae* to selected agents.

## EXPERIMENTAL PROCEDURES

### *In-silco analysis of gene expression omnibus profile database* (GEO)

The Gene Expression Omnibus (GEO) profile database (https://www.ncbi.nlm.nih.gov/geoprofiles/) was searched using gene symbol (SAF1) to study the expression profiles during stress in *S. cerevisiae* wild type cell. The profile data was studied for expression status of SAF1 gene in WT cells, treated with stress causing agent.

### *In-silico analysis of transcriptional regulation databases* (YEASTRACT)

The YEASTRACT database (http://www.yeastract.com) of *Saccharomyces cerevisiae* was searched for association between SAF1 and AAH1 gene and global transcription factor during normal and stress condition. Further search was carried out for regulatory association of SAF1 and AAH1 with the transcription factors during stress condition and ranked for given TF under the stress condition.

### In-silco analysis of yeast stress expression database (yStreX)

The Yeast stress expression database (http://www.ystrexdb.com/) was analysed for expression of SAF1, AAH1, and RLM1 transcription factor. An advanced search was conducted for SAF1 and RLM1 genes at default cut-off (log2 Fold Change > 1.5) and (adjusted p-value at < 0.05) under all experimental class. The results on the expression status of genes included the statistical values and biological features for specific conditions.

### Yeast Strains and plasmids

*Saccharomyces cerevisiae* strain, BY4741 (*Mata his3Δ1 leu2Δ0 met15Δ0 ura3Δ0)* and JC2326 (*MAT-ura3, cir0, ura3–167, leu: his, his32 <u>Ty1his3AI-270</u>, Ty1NEO-588, Ty1 (tub: lacs)-146)* were used in this study (**Table 1)**. The list of plasmids used for generation of marker cassette mentioned in the **Table 2** and primers in **Table 3**

**Table 1:** Yeast strains and their genotype used in this study

| S.no. | Strain | Genotype | Source |
| --- | --- | --- | --- |
| 1 | BY4741 (WT) | *MATa his3Δ1 leu2Δ0 met15Δ0 ura3Δ0* | Dr. Deepak Sharma IMTECH |
| 2 | *MSR1* | *saf1Δ::HIS3MX* | This study |
| 3 | *MSR2* | *rlm1Δ::KanMX* | This study |
| 4 | *MSR3* | *saf1Δ::HIS3MX, rlm1Δ::KanMX* | This study |

**Table 2:**
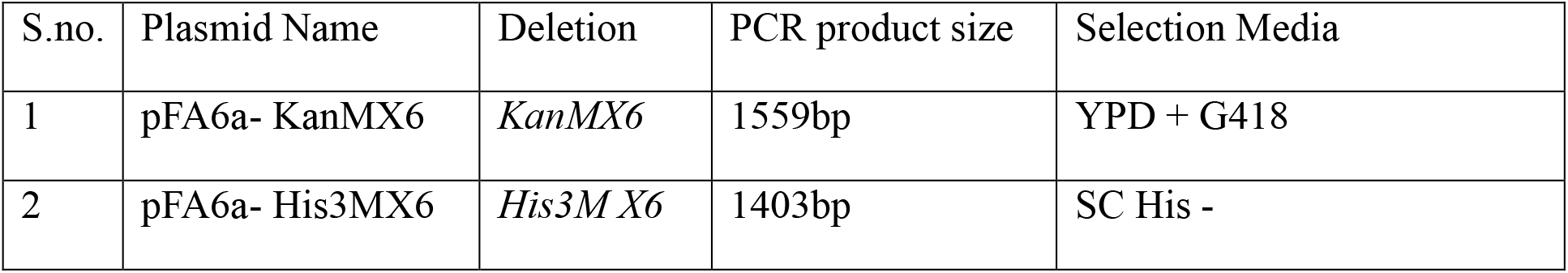
List of plasmids used for generating deletion cassette

| S.no. | Plasmid Name | Deletion | PCR product size | Selection Media |
| --- | --- | --- | --- | --- |
| 1 | pFA6a- KanMX6 | *KanMX6* | 1559bp | YPD + G418 |
| 2 | pFA6a- His3MX6 | *His3MX6* | 1403bp | SC His - |

**Table 3:**
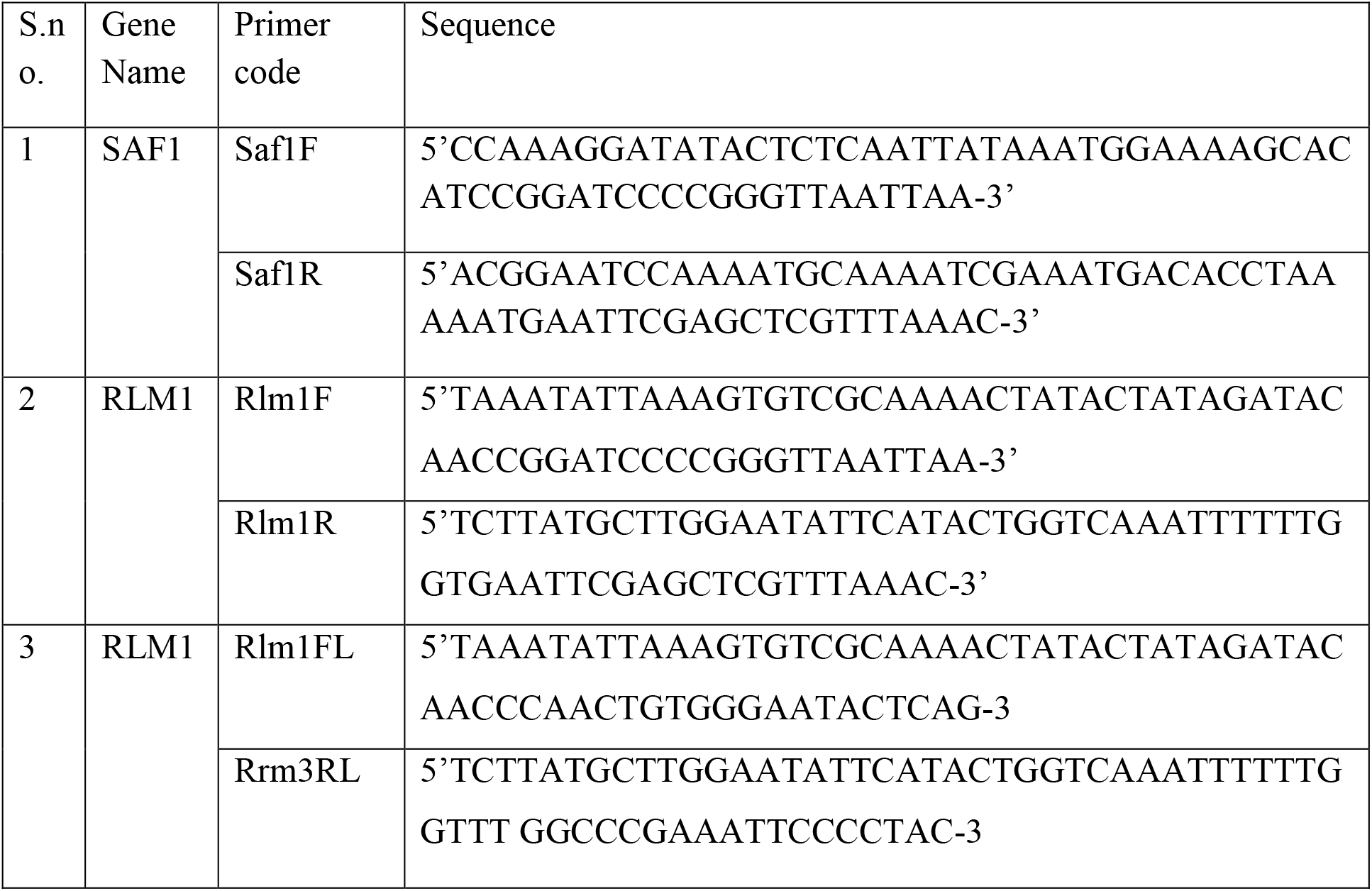
List of primers used for construction of deletion strains

### Growth Assay

For growth assessment of the WT (BY4741), *saf1Δ*, *rlm1Δ* and *saf1Δrlm1Δ* in YPD broth, optical density (OD600nm) measurement was carried using TOSHVIN UV-800 SHIMADZU spectrophotometer. The growth of strains was measured at 2 hrs of interval for 14 hrs, three independent replicates of each culture’s OD were measured, and average of that was plotted against the time. Growth of the strains was also compared by streaking on the YPD plates followed by incubation at 30°C for 2-3 days.

### Phase Contrast Microscopy

To compare the morphology of WT and deletion strain, phase contrast microscopy was carried out on log phase grown strains in YPD medium at 30°C using Leica DM3000 microscope at 100X magnification.

### Calcofluor white staining and Fluorescence imaging

For staining of wild type and mutant cells (WT, *saf1Δ, rlm1Δ,* and *saf1Δrlm1Δ)* with Calcofluor white stain method mentioned (Sharma et. al 2019, biorxiv archived data www.biorxiv.org) was adopted. Briefly, WT and each mutant strain were grown over night at 30°C and next day were re-inoculated in fresh YPD medium in 1:10 ratio. Cells were grown to log phase and collect by centrifugation. The collected cells were suspended in 100μl of solution containing Calcofluor white (50 μg/ml solution) fluorescent dye. Cells observed under 100X magnification using Leica DM3000 fluorescence microscope using Ultraviolet filters as per instructions mentioned in the handbook.

### Spot Assay

The spotting assay was performed as mentioned in the (Sharma et. al 2019, biorxiv archived data www.biorxiv.org) for comparative assessment of the growth fitness among the WT and mutant strains in presence of stress causing agents such as Hydroxyurea (HU), Methyl methane sulfonate (MMS), Nocodazole, Benomyl, Calcofluor White, SDS, NaCl, Glycerol, DMSO, H_2_O_2_ . Briefly, wild type (BY4741) and its deletion derivatives strains (*saf1Δ, rlm1Δ,* and *saf1Δrlm1Δ*) were grown in the 25 ml YPD (Yeast Extract 1% w/v, Peptone 2% w/v, dextrose 2% w/v) medium overnight at 30°C. The next day the cultures were diluted and grown in fresh YPD medium for 3-4 hrs so as to reach log phase (OD600 0.8-1.0). The cultures concentration was adjusted by dilution to that they are equal by OD at 600nm followed by ten-fold serial dilution and spotting of 3μl volume from each dilution onto YPD agar plates with and without stress agents. The plates were incubated at 30°C for 2-3 days and cellular growth of the WT and mutants were recorded.

### Assay for Ty1 retromobility

The Ty1 retro-mobility assay was performed as per method mentioned in the study (SCHOLES *et al.* 2001; BAIRWA *et al.* 2011) Sharma et. al 2019, biorxiv archived data www.biorxiv.org.). The reporter strain JC2326 was used for construction of deletion derivatives (*saf1Δ, rlm1Δ,* and *saf1Δrlm1*Δ). Each strain carried a single Ty1 element marked with indicator gene HIS3AI. The indicator gene HIS3 is interrupted by artificial intron (AI) in an orientation opposite to the HIS3 gene transcription. During transcription of marked Ty1 with this arrangement followed by splicing and reverse transcription, generates the cDNA with functional HIS3 ORF which upon integration into genomic locations generate HIS3 positive phototrophs. The quantitative measurement of the HIS3 positive colonies was considered as frequency of retro-mobility. To measure the Ty1 retro-transposition frequency in the WT (JC2326; reporter strain) and the deletion derivatives (*saf1Δ rlm1Δ* and *saf1Δrlm1Δ*), a single colony of each strains was inoculated into 10 ml YPD broth and grown overnight at 30°C.The overnight grown cultures were again inoculated in 5 ml YPD at 1:1000 dilutions. The cultures were grown for 144 hrs at 20°C. The culture was serially diluted and plated on minimal media (SD/His^−^ plates) followed by incubation at 30°C for 3-7 days. The three independent replicates for each strain was used for serial dilution and plated on SD/His^−^ plates. The numbers of HIS+ colonies were counted from each plate and plotted for comparative assessment of Ty1 frequency.

### Assay for Nuclear Migration defects using Fluorescence microscopy

For detection of nuclear migration defects, assay described by (PALMER *et al.* 1992; BRACHAT *et al.* 1998) and (Sharma et. al 2019, biorxiv archived data; www.biorxiv.org) was adopted. The number of nuclei per cell in yeast strains was determined with nuclear binding dye (DAPI). Briefly, strains were grown to early log phase (OD_600_ ~ 0.8) at 30ºC. Yeast cells were washed with distilled water and suspended in 1X PBS (Phosphate Buffer Saline). Further, fixation was done by addition of 70% ethanol before DAPI staining. Cells were washed with 1X PBS buffer then again centrifuged for 1 minute at 2500 rpm. DAPI stain (1mg/ml stock) to final concentration of 2.5μg/ml was added and incubated for 5 minutes at room temperature and visualized under fluorescent microscope with 100X magnification. A total of 200 cells were counted and grouped according 0, 1, 2, and multi nuclei per cell, more than two nuclei per cell indicated the nuclear migration defect.

### Statistical methods

Statistical significance of observations was determined using paired student t-test. P-value less than 0.05 indicated significant.

## Results

### SAF1 overexpressed in WT cells treated with stress agents

The GEO profile analysis showed elevated expression of SAF1 gene during, Clioquinol, Pterostilbene, Gentamicin, hypoxia, genotoxic, desiccation, and heat stress in WT cells (**Figure S1**–**S7** and **Table 4**; source GEO profile data base NCBI).

**Table 4:** Gene Expression Omnibus profile of the SAF1 gene during stress (source: GEO)

| S.no. | Stress Agent | Upregulation of SAF1 WT cells during Stress | GEO Profile ID |
| --- | --- | --- | --- |
| 1 | Clioquinol | ↑ | ID: 66726869 |
| 2 | Pterostilbene | ↑ | ID: 52693374 |
| 3 | Gentamicin | ↑ | ID: 46082374 |
| 4 | Hypoxia | ↑ | ID: 57031974 |
| 5 | Genotoxic | ↑ | ID: 11783774 |
| 6 | Desiccation | ↑ | ID: 38619274 |
| 7 | Heat Shock | ↑ | ID: 89011 |

### RLM1 transcription factor associates with SAF1 expression during stress

An in-silico **a**nalysis of YEASTRACT database (http://www.yeastract.com) of *Saccharomyces cerevisiae* for association of global transcription factors with SAF1 during normal and stress condition revealed a total of three transcription factors (Rlm1, Spt23 and Cin5) (**Table 5**), among three Rlm1 and Cin5 found to be positive regulator. However six transcriptions factors (Rpn4p, Msn2p, Msn4p, Aft1p, Rap1p, and Rlm1p) showed association with AAH1 during stress (**Table 5)**. Rlm1 found to be common transcription regulator both SAF1 and AAH1 genes (**Figure 1**). A regulatory association analysis showed RLM1 as positive regulator of the SAF1 gene and negative regulator of the AAH1 gene. Furthermore, analysis of yeast stress expression database (yStreX) showed 2-fold elevated expression of SAF1 and RLM1 upon stress condition (**Table 6**). Based on *in-silico* analysis we suggest that upon exposure to stress, expression of RLM1 leads to upregulation of SAF1 and downregulation of AAH1, resulting in quiescence phase induction during stress, due to reduction in Aah1 activity both transcriptionally and post-transcriptionally. Further, we hypothesized that deletion of SAF1 and RLM1 together may contribute to stress tolerance.

**Table 5:**
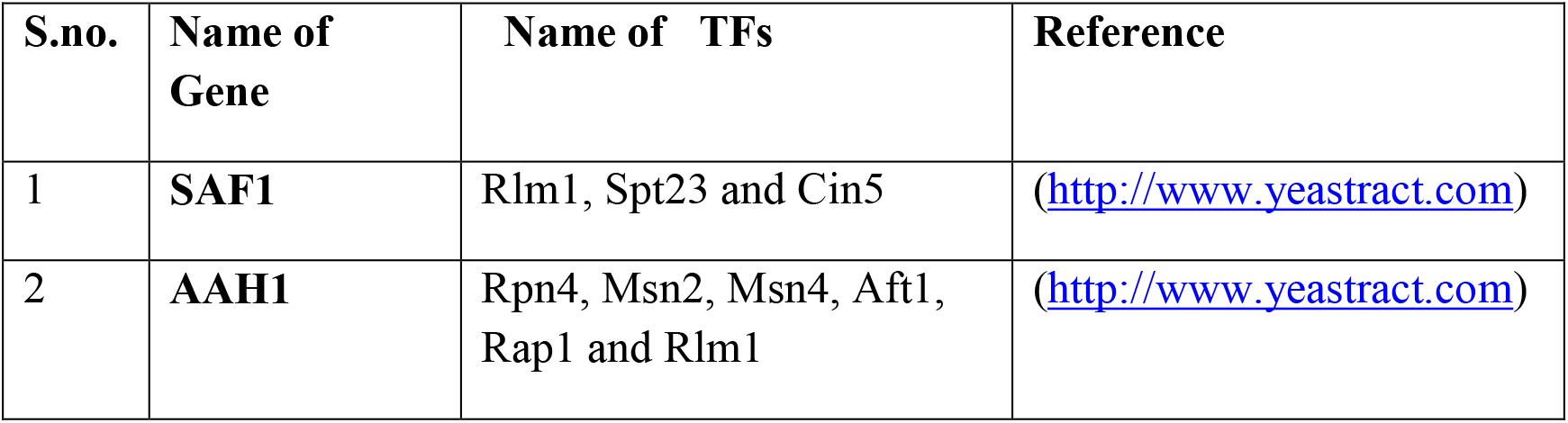
List of global transcription factors regulating SAF1 and AAH1 during normal and stress condition (Source: YEASTRACT database)

**Table 6:** Expression of SAF1 and RLM1 during stress condition (source: yStreX (http://www.ystrexdb.com)

| <u>S.no</u> | <u>Gene</u> | <u>Class</u> | <u>Sub- class</u> | <u>Contro l</u> | <u>Case</u> | <u>Strain</u> | <u>Log FC</u> | <u>P value</u> |
| --- | --- | --- | --- | --- | --- | --- | --- | --- |
| 1 | **RLM1** | Cultivation | Fermentation | 2 day | 14 days | 285 | **2.282** | 0 |
| 2 | **SAF1** | Extra stress | Desiccation | Control | 100% | BY4743 | **2.142** | 6.1E-13 |
| 3 | **SAF1** | Limiting | Amino acid (AA- Lim) | YPD | 15 min and SD media | W303a | **2.224** | 5.4E-11 |
| 4 | **SAF1** | Media | SD | YPD | 15 min and SD media | BY4741 | **2.224** | 5.4E-11 |
| 5 | **SAF1** | Inhibitor | Rapamycin | Control | 200ng/ml | BY4741 | **2.057** | 9.91154747E-6 |
| 6 | **SAF1** | Temperature | 37°C | 25°C | 5 min | BY4741 | **2.006** | 4.88851412E-6 |
| 7 | **SAF1** | Inorganic | Hydrogen peroxide | Control | 0.4mM | BY4741 | **2.185** | 8.132709E-8 |

**Figure 1:**
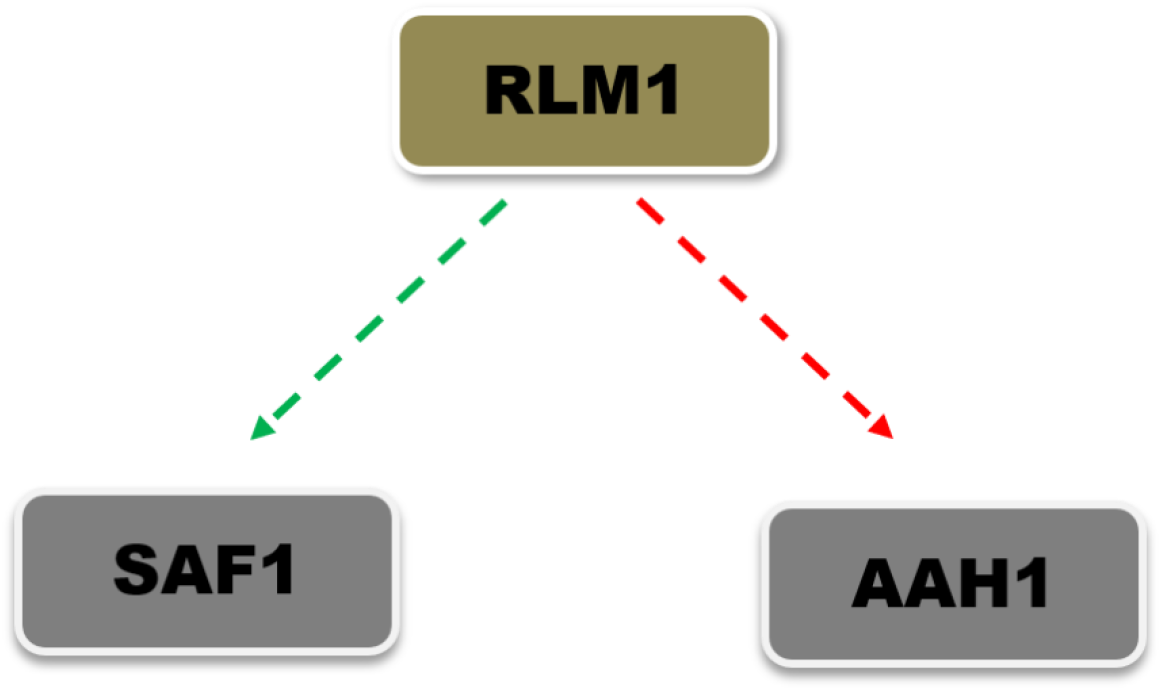
Rlm1 as novel transcription factor which regulates the SAF1 and AAH1 expression during stress. YEASTRACT database analysis showed that Rlm1 as common regulator of both the genes, where it showed positive association with SAF1 whereas negative association with AAH1.

### Loss of SAF1 and RLM1 together showed no growth defects

Comparative growth assessment of WT, *saf1Δ, rlm1Δ and saf1Δrlm1Δ* in liquid and solid rich medium showed growth reduction in *saf1Δ, rlm1Δ and saf1Δrlm1Δ* as compared to WT (**Figure 2, A C)**. Similarly, no distinguishable difference was observed in the morphology of the WT and mutant strain (**Figure 2 B)**.

**Figure 2:**
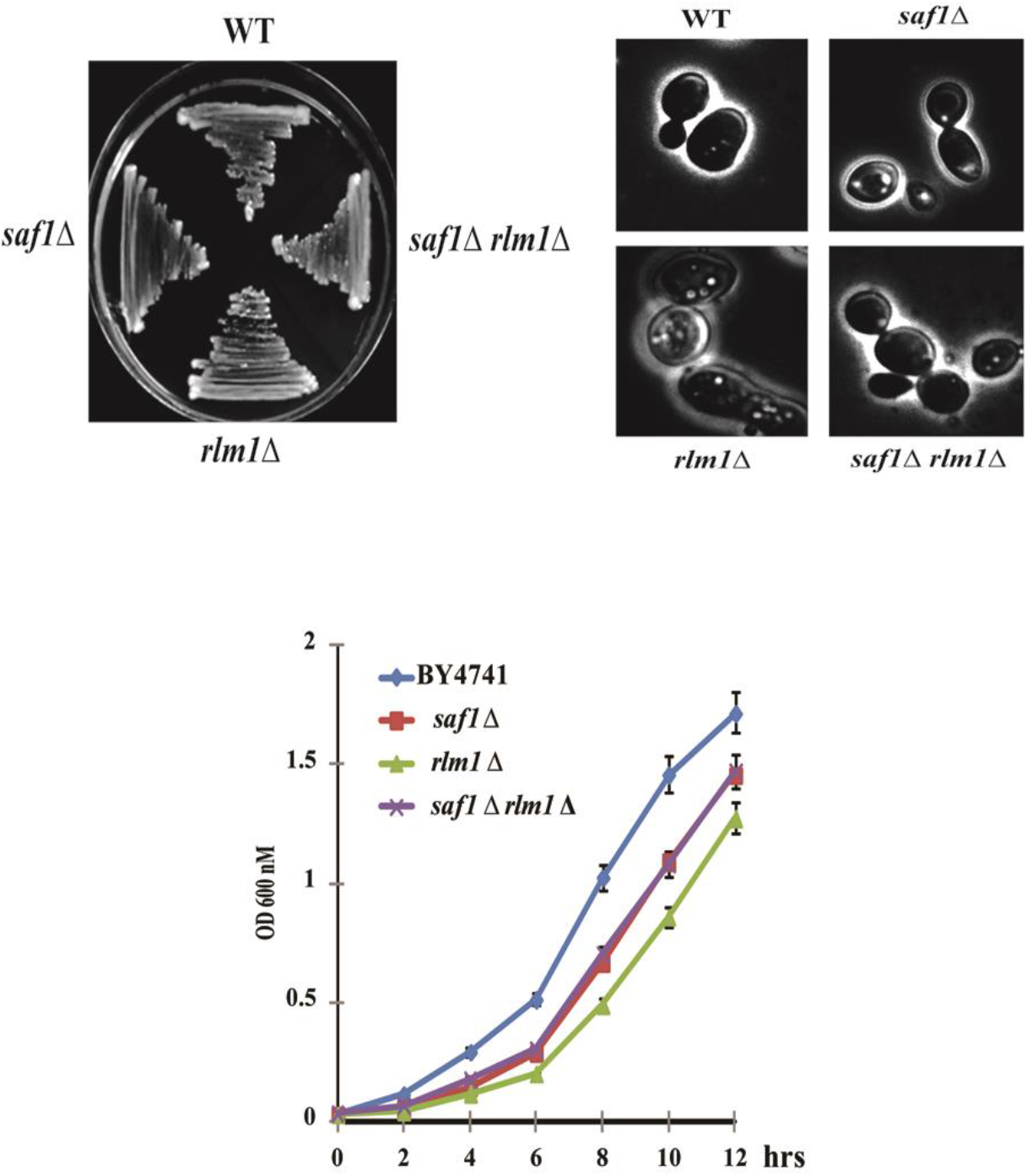
Comparative analysis of growth and morphology of WT,*saf1Δ, rlm1Δ, saf1Δrlm1Δ* cells. **A.** Growth of streaked strains on YPD agar plates incubated for 2 days at 30°C and photographed. **B.** Phase contrast images of log phase cultures at 100X magnification using Leica DM3000, no major difference with regard to morphology observed with mutants and WT **C.** Growth kinetics of WT, *saf1Δ, rlm1Δ, saf1Δrlm1Δ.* Both single and double mutant showed slightly slow growth in comparison to WT. Cells were collected every 2 hour period and cellular growth was measured by optical density (OD) at 600 nm using TOSHVIN UV-1800 SHIMADZU. The data shown represent the average of three independent experiments. The error bars seen represent the standard deviation for each set of data.

### Loss of SAF1 and RLM1 together leads to tolerance to Calcofluor White and SDS stress

Calcofluor white is a fluorochrome stain which perturbs the cell wall whereas SDS is membrane disrupting agent. Calcofluor binds to chitin and cellulose. The Calcofluor white stained cell observed under fluorescence microscope showed mother daughter neck region stained in WT *S. cerevisiae*. The *saf1Δrlm1Δ* stained cell showed staining of entire cell wall region when compared with WT and s*af1Δ* (**Figure 3 A**). The relative fluorescence intensity was also synergistically elevated in the s*af1Δrlm1Δ* cell in comparison to either s*af1Δ* or *rlm1Δ (***Figure 3 B**). The cellular growth response of WT, s*af1Δ, rlm1Δ* and s*af1Δrlm1Δ* strains in calcofluor white (30μg/ml) and SDS (0.0075%) showed tolerance (**Figure 3 C, D).**

**Figure 3:**
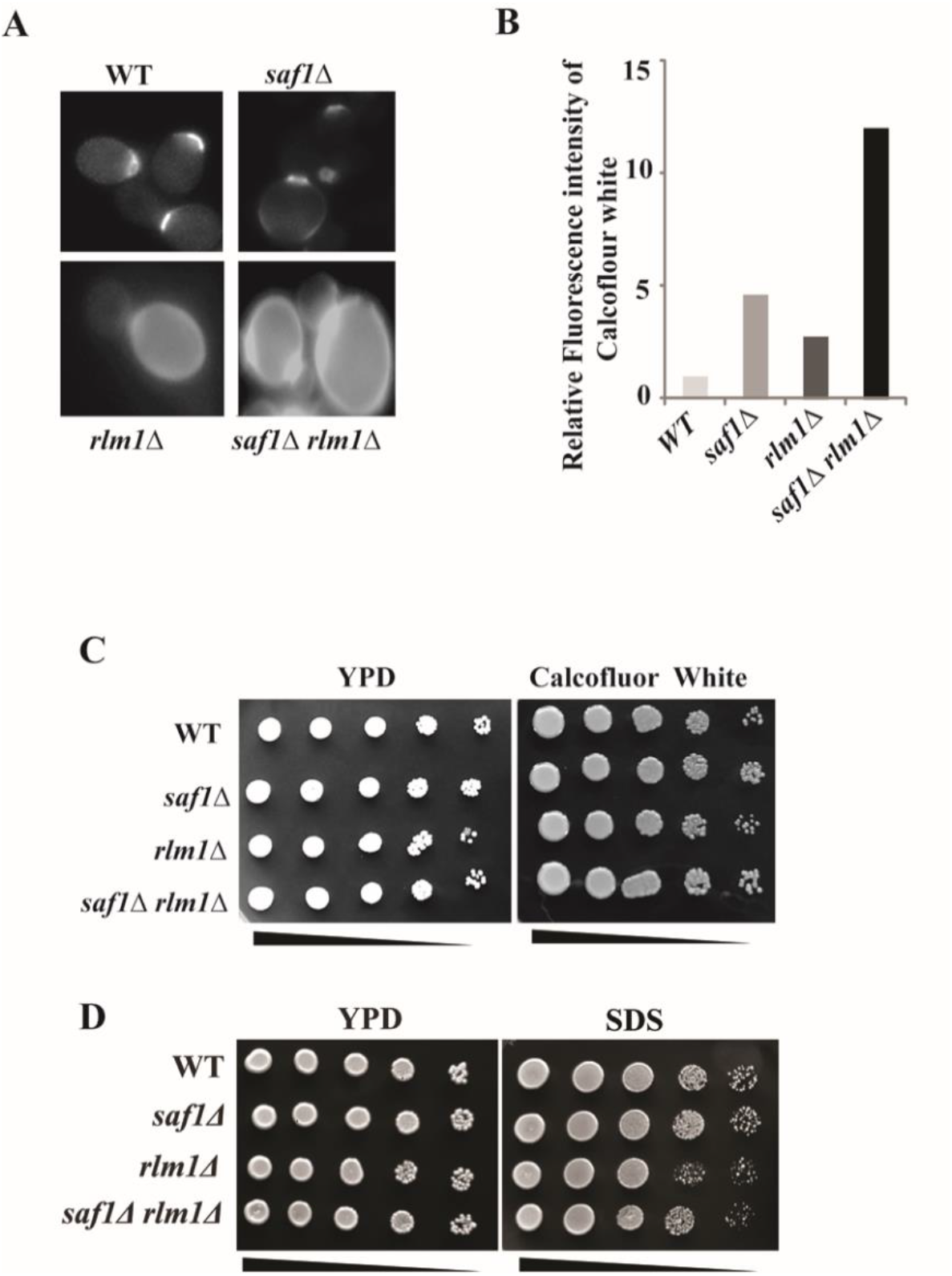
Loss of SAF1 and RLM1 together showed Stress tolerance to Calcofluor White, SDS and altered chitin distribution. **A.** Calcofluor stained WT, *saf1Δ, rlm1Δ, saf1Δrlm1Δ* cells, which is specific to chitin in the cell wall. The *rlm1Δ* and *saf1Δrlm1Δ* showed the altered chitin in the cell wall in comparison to WT and *saf1Δ* cells **B.** Relative fluorescence intensity of the Calcofluor white stained cells of s*af1Δrlm1Δ* showed 10-fold increases in the intensity in comparison to WT cells. **C.** Cellular growth response of WT, *saf1Δ, rlm1Δ,* and *saf1Δrlm1Δ* cells in presence of 30μg/ml Calcofluor white. Single and double mutant showed the tolerance to CFW as similar to WT. **D.** Cellular growth response of WT, *saf1Δ, rlm1Δ, saf1Δrlm1Δ* cells in presence of 0.0075% SDS, which is membrane disrupter. The *saf1Δrlm1Δ* showed slight sensitivity to SDS in comparison to *saf1Δ, rlm1Δ* and WT.

### Loss of SAF1 and RLM1 together leads to tolerance to H_2_O_2_ and DMSO

Hydrogen peroxide (H_2_O_2_) and Dimethyl sulfoxide (DMSO) both are oxidative stress causing agents (KWAK *et al.* 2010; SADOWSKA-BARTOSZ *et al.* 2013; QI *et al.* 2019). We investigated the cellular growth response of the WT and *saf1Δ, rlm1Δ,* s*af1Δrlm1Δ* cells in presence of 2mM H_2_O_2_ and 8% DMSO using spot assay. We observed that WT, s*af1Δ, rlm1Δ,* and s*af1Δrlm1Δ* cells showed the tolerance to the indicated concentration of H_2_O_2_ and DMSO (**Figure 4A, B).**

**Figure 4:**
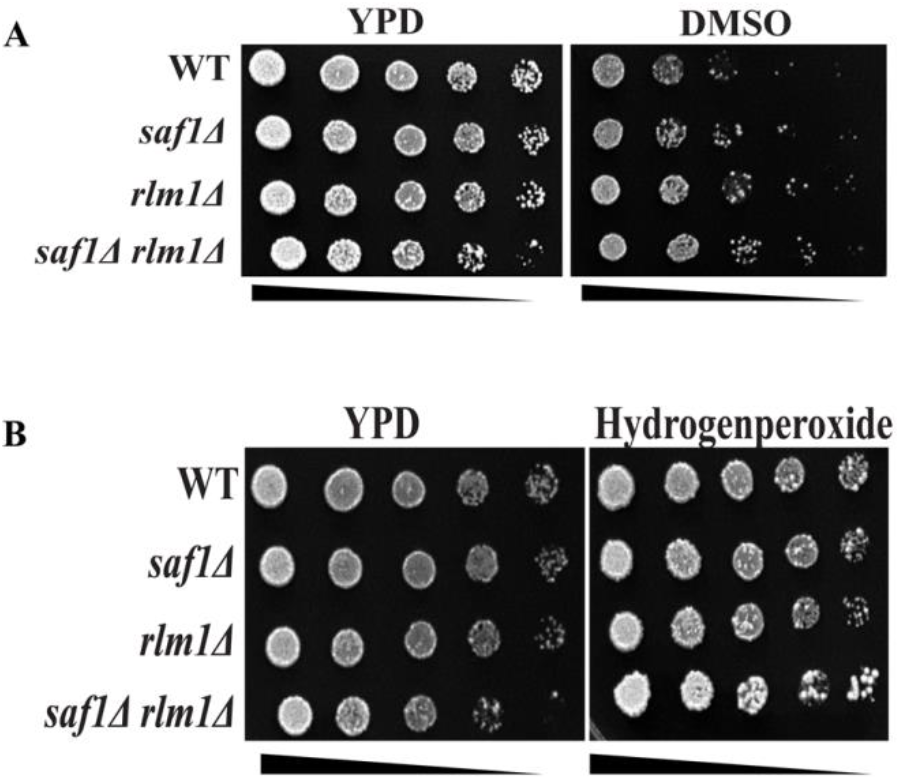
Loss of SAF1 and RLM1 together showed DMSO and H_2_O_2_ tolerance in comparison to WT cells. **A.** Cellular growth response of WT, *saf1Δ, rlm1Δ, and saf1Δrlm1Δ* cells in presence of 8% DMSO. The *saf1Δrlm1Δ* cells showed tolerance to DMSO in comparison to WT **B.** Cellular growth response of WT, *saf1Δ, rlm1Δ, saf1Δrlm1Δ* cells in presence of 2mM H_2_O_2_. The *saf1Δrlm1Δ* cells showed tolerance to H_2_O_2_ in comparison to WT.

### Loss of SAF1 and RLM1 leads to tolerance to Glycerol and NaCl stress

A high concentration of external glycerol and NaCl leads to expression of osmotic stress response genes. We tested cellular growth response of WT and *saf1Δ, rlm1Δ,* and *saf1Δrlm1Δ* cells in 4% of glycerol and 1M NaCl. We observed that WT, *saf1Δ, rlm1Δ,* and *saf1Δrlm1Δ* showed tolerance to 4% glycerol and 1M NaCl in spot assay. (**Figure 5A, 5B)** whereas failed to grow in presence of 1.4M NaCl (data not shown). This suggests that *saf1Δrlm1Δ* cell tolerates osmotic stress mediated by glycerol and NaCl.

**Figure 5:**
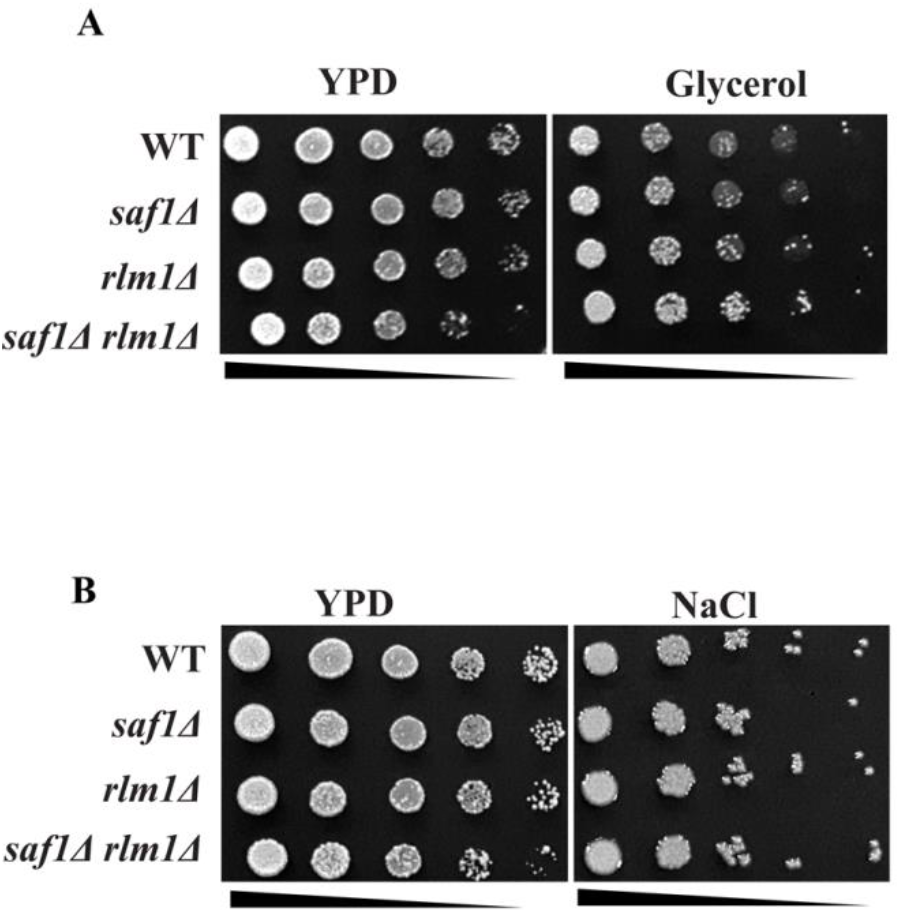
Loss of SAF1 and RLM1 contributes to Glycerol and NaCl tolerance. **A.** Cellular growth response of WT, *saf1Δ, rlm1Δ, and saf1Δrlm1Δ* cells in presence of 4% glycerol. The *saf1Δrlm1Δ* cells showed tolerance to glycerol in comparison to WT **B.** Cellular growth response of WT, *saf1Δ, rlm1Δ, saf1Δrlm1Δ* cells in presence of 1M NaCl. The *saf1Δrlm1Δ* cells showed tolerance to 1M NaCl similar to WT.

### Loss of SAF1 and RLM1 together leads to sensitivity to genotoxic stress

DNA alkylating agent, methyl methane sulfonate (MMS) elicit DNA damage in cells, which results in activation of DNA damage repair pathways. Hydroxyurea acts as replication checkpoint inhibitor and causes DNA replication defects by inhibiting the activity of ribonucleotide reductase (RNR). Here, we tested the cellular growth response of the WT, s*af1Δ, rlm1Δ,* and s*af1Δrlm1Δ* cells in presence of 0.035% MMS and 200mM HU by spot assay. We observed that s*af1Δrlm1Δ* cells showed growth pattern similar to WT, s*af1Δ* and *rlm1Δ* cells however sensitive to hydroxyurea in comparison to WT, s*af1Δ and rlm1Δ* cells (**Figure 6B, C**).

**Figure 6:**
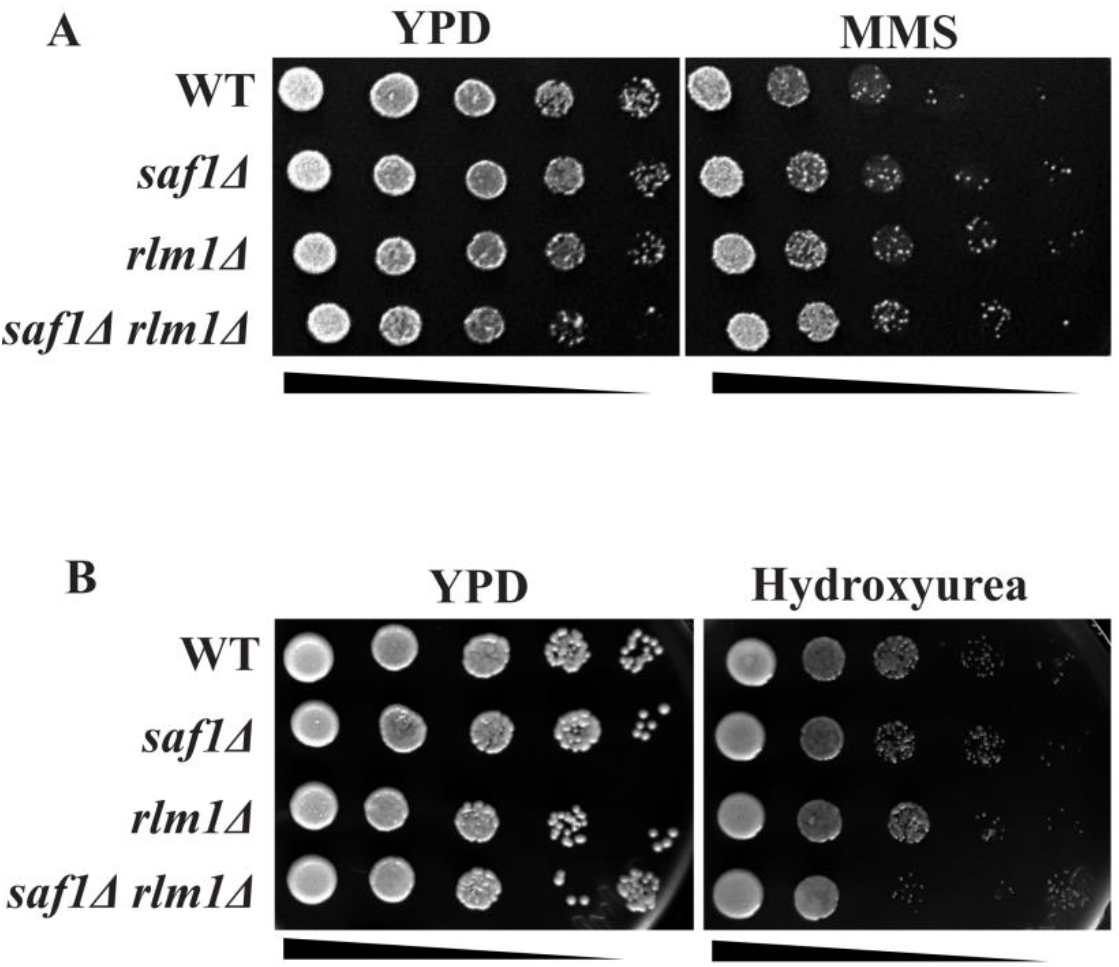
Loss of SAF1 and RLM1 together showed sensitivity to genotoxic stress agents MMS and HU. **A.** Cellular growth response of WT, *saf1Δ, rlm1Δ, saf1Δrlm1Δ* cells in presence of 0.035% MMS. The *saf1Δrlm1Δ* cells showed sensitivity to MMS in comparison to WT. **B.** Cellular growth response of WT, *saf1Δ, rlm1Δ, saf1Δrlm1Δ* cells in presence of 200mM HU. The *saf1Δrlm1Δ* cells showed sensitivity to HU in comparison to WT, *saf1Δ* and *rlm1Δ* cells.

### Loss of SAF1 and RLM1 together leads to sensitivity to Benomyl and Nocodazole

Benomyl and Nocodazole both are microtubule depolymerizing agents which lead to apoptosis (THOMAS et al. 1985; JORDAN and WILSON 2004). We investigated the cellular growth response of WT *saf1Δ, rlm1Δ, and saf1Δrlm1Δ* cells in presence of 100μg/ml benomyl and 50μg/ml Nocodazole. We observed that WT, *saf1Δ, rlm1Δ* and *saf1Δrlm1Δ* showed sensitivity to both microtubule drugs in spot assay (**Figure 7A, 7B**). The image analysis of the cells exposed to Hydroxyurea and Nocodazole showed characteristic elongated buds in WT, *saf1Δ, rlm1Δ,* and *saf1Δrlm1Δ* as compare to the cells grown on rich media without HU and Nocodazole (**Figure 8**).

**Figure 7:**
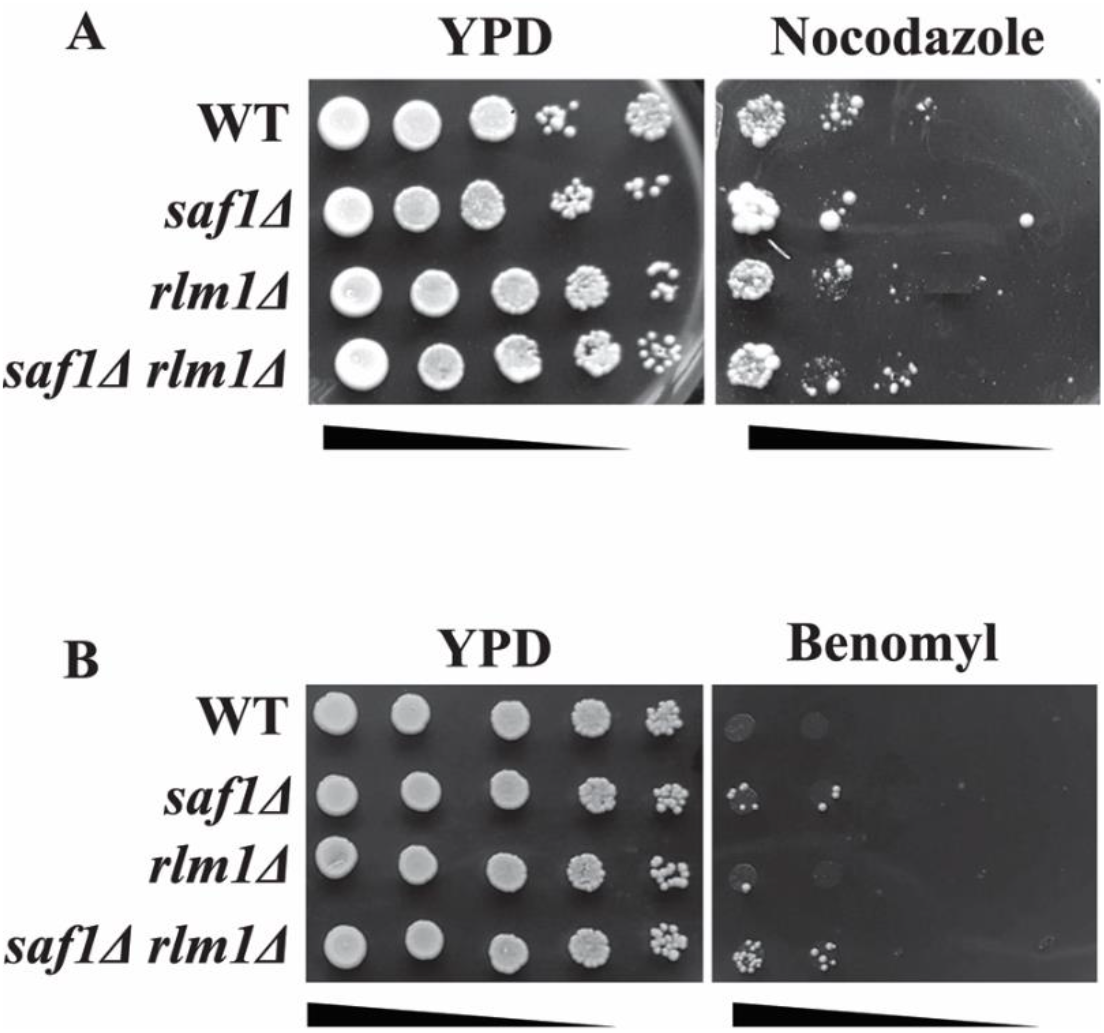
Loss of SAF1 and RLM1 together showed sensitivity to Nocodazole and Benomyl. **A.** Cellular growth response of WT, *saf1Δ, rlm1Δ, saf1Δrlm1Δ* cells in presence of 50μg/ml Nocodazole. The *saf1Δrlm1Δ* cells showed sensitivity to Nocodazole when compared with growth on YPD. **B.** Cellular growth response of WT, *saf1Δ, rlm1Δ, saf1Δrlm1Δ* cells in presence of 100μg/ml benomyl. The *saf1Δrlm1Δ* cells showed sensitivity to benomyl when compared with growth on YPD.

**Figure 8:**
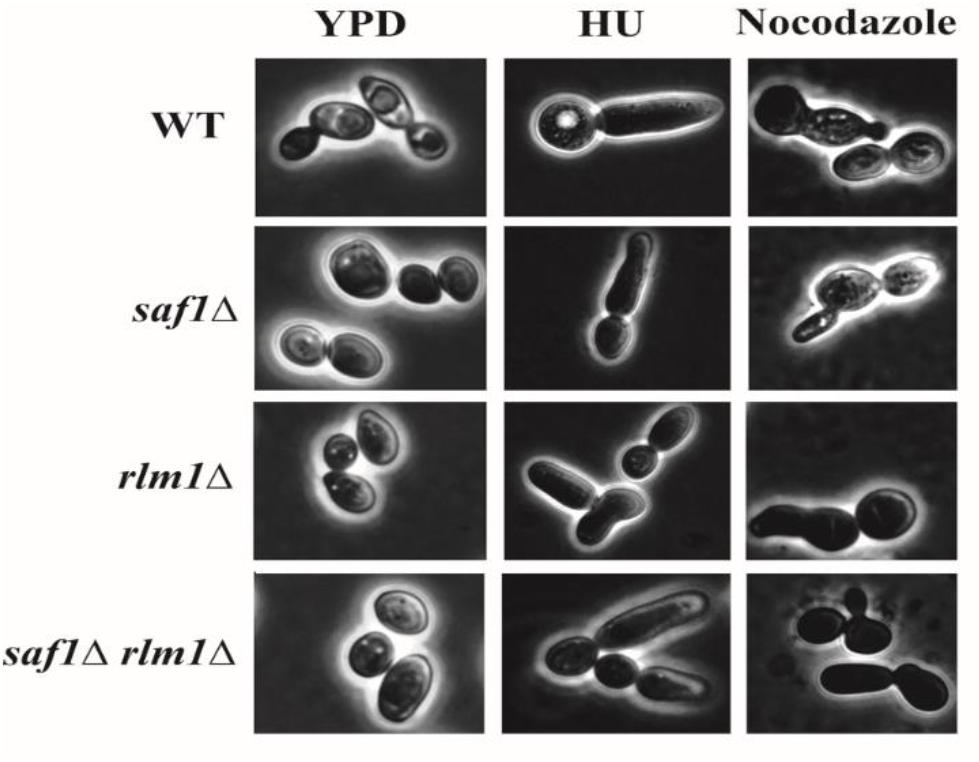
Comparative morphological feature of WT,*saf1Δ, rlm1Δ, saf1Δrlm1Δ* cells grown on YPD media and YPD media with HU and Nocodazole. The WT, *saf1Δ, rlm1Δ,* and *saf1Δrlm1Δ* showed the growth defects due inhibition of cell division by both HU and Nocodazole.

### Loss of SAF1 and RLM1 together showed increased multi-nuclei phenotype

The absence of both the genes leads to sensitivity to genotoxic stress, suggesting there may be defect in nuclear DNA division. We investigated the status of nuclear DNA in the cells using DAPI in WT, *saf1Δ, rlm1Δ,* and s*af1Δrlm1Δ* cells (**Figure 9 A)**. We observed that 80% WT cells, showed compact nuclei as single dot whereas in 20% *saf1Δ* and 28% *saf1Δrlm1Δ* cells showed the multi nuclei phenotype (**Figure 9 B)**. However, multi-nuclei phenotype was not observed in *rlm1Δ* cells.

**Figure 9:**
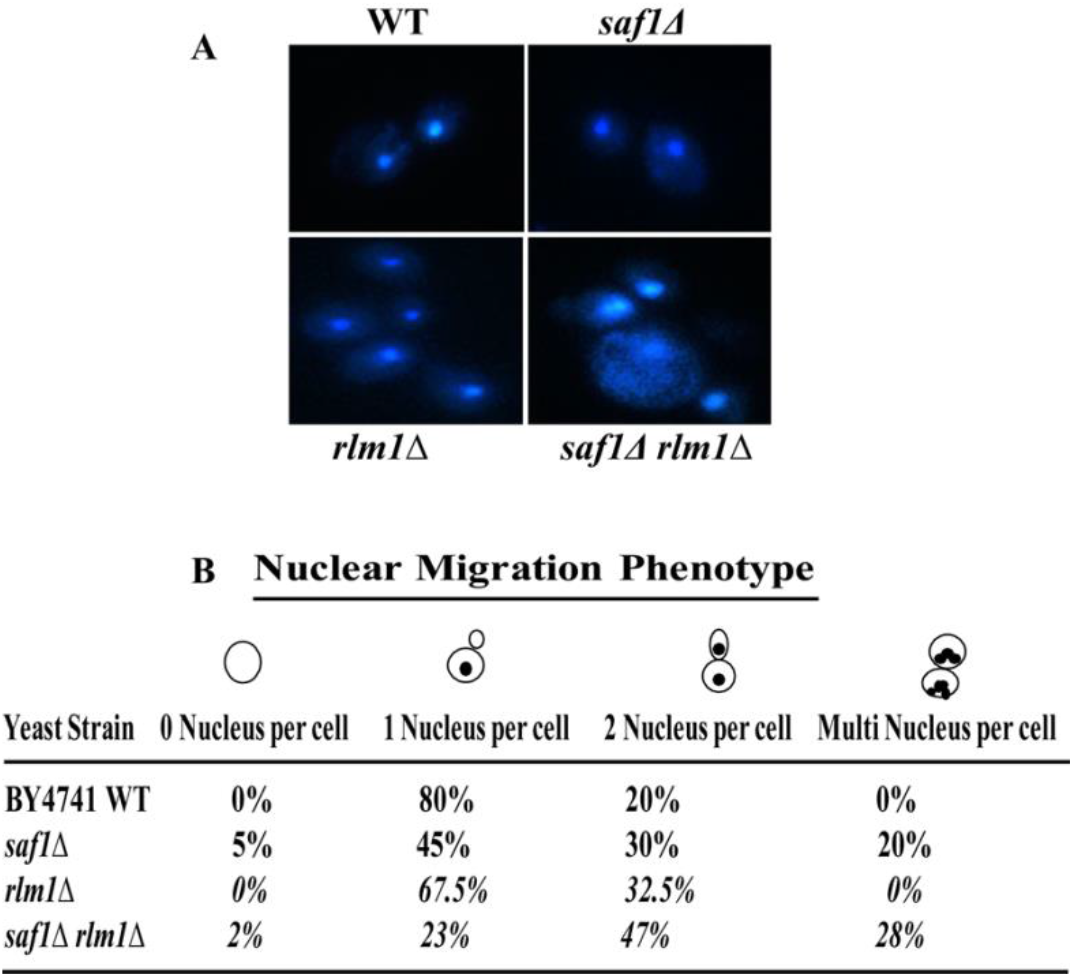
Loss of SAF1 and RLM1 together showed increased multi-nuclei phenotype. Assessment of nuclear migration phenotype using DAPI (4′,6-diamidino-2-phenylindole) staining of WT, *saf1Δ, rlm1Δ, saf1Δrlm1Δ* A. Representative fluorescent images WT, *saf1Δ, rlm1Δ, saf1Δrlm1Δ* showing the status of Nuclear DNA, images were acquired at the 100X magnification using Leica DM3000 fluorescent microscope. **B** Table showing the percentage from the count of 200 cells as, 0, 1, 2 and multi- nucleus in each strain, more than two nuclei indicate the nuclear migration defect.

### Loss of SAF1 and RLM1 together showed elevated Ty1 retro-transposition

DNA replication stress and DNA damage both leads to increase in Ty1 retro-mobility in *S. cerevisiae.* We studied the Ty1 retromobility in the *WT*, *rlm1Δ, saf1Δ* and *saf1Δrlm1Δ* genetic background. We observed ~ 40-fold increase in the Ty1 retromobility in the *rlm1Δ* whereas *saf1Δrlm1Δ* cells showed ~120-fold increase in Ty1retromobility in comparison to WT (**Figure 10 A, B**).

**Figure 10:**
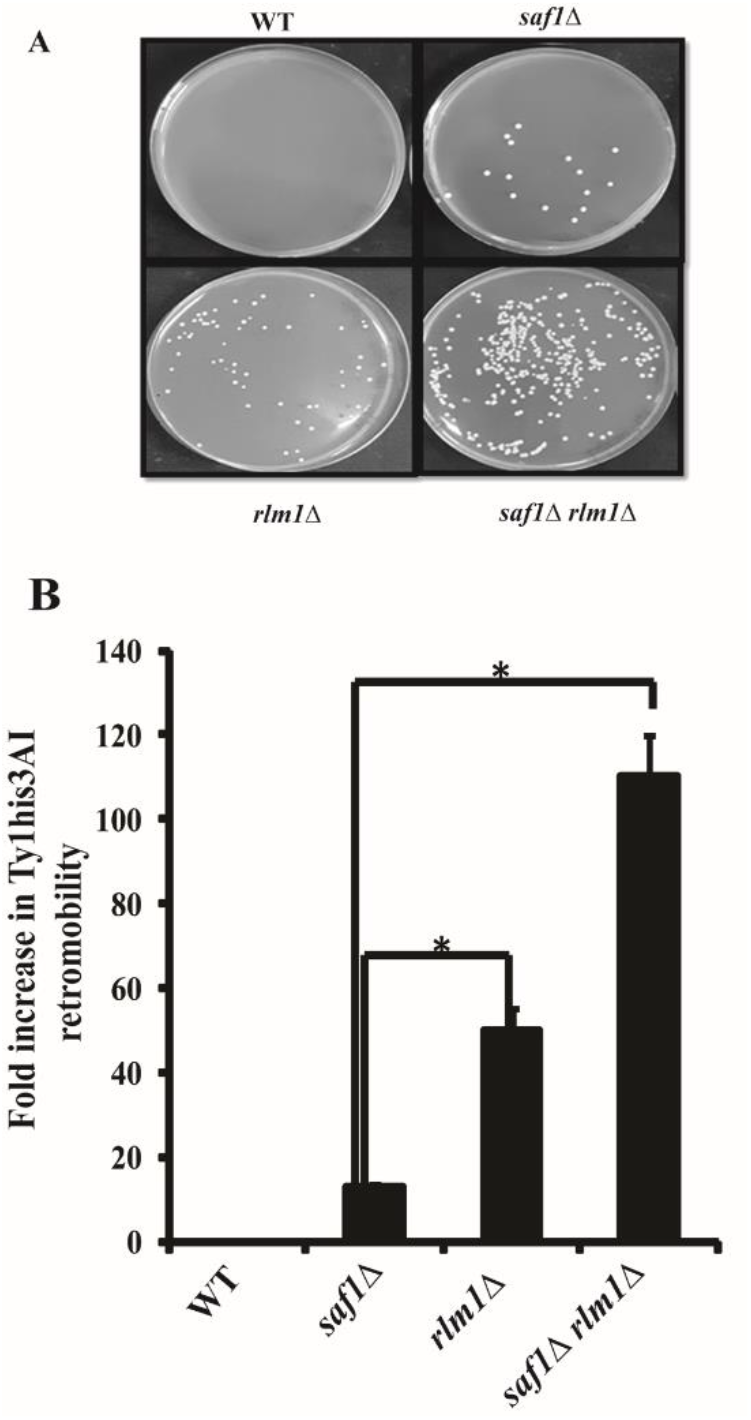
Loss of SAF1 and RLM1 together showed elevated Ty1 retro-transposition. Assay for HIS3AI marked Ty1 transposition frequency in WT, *saf1Δ, rlm1Δ, saf1Δrlm1Δ* **A.** Images of plates showing the Ty1 transposition induced colonies on SD plate lacking His media. **B** Bar diagram showing the frequency of Ty1his3AI transposition in each strain. The data shown represent the average of three independent experiments. The significance of retrotransposition was determined using two tailed t-test. P -value 0.05 indicate significant difference.

## Discussion

Here in this study we report the genetic interaction between transcription factor RLM1 and F-box motif encoding gene SAF1. The expression of SAF1 was elevated during various stress conditions in WT cells as indicated by GEO profile analysis. These initial observations lead us to search for transcription factor may be associated with expression of SAF1 during stress. The search for transcription factor using YEASTRACT database suggested Rlm1 as novel transcription factor for SAF1 during stress. The regulatory association search between SAF1, AAH1 and RLM1 showed that RLM1as positive regulator of SAF1 and negative regulator of AAH1 genes expression. Further analysis using yStreX database suggested that both SAF1 and RLM1 expression was elevated during the stress condition in WT cells whereas the AAH1 expression was downregulated. These observations supported the AAH1 gene model, where during phase transition from proliferative state to quiescence phase upon nutrition stress, Saf1 recruits Aah1 for ubiquitination and degradation. It is expected that during stress condition the transcription of AAH1 should be downregulated and SAF1 upregulated. The RLM1 involvement was not discovered. We report that RLM1 as transcription factor regulates the expression of SAF1 and AAH1 during stress. Based on *in-silico* analysis, it was hypothesized that ablation of SAF1 and RLM1 together may lead to stress tolerance in the double mutant. Our experimental analysis validated the hypothesis, where double mutant cells showed the resistance to Calcofluor white, SDS and hydrogen peroxide. Calcoflour white and SDS both are cell wall and cell membrane disruptor whereas H_2_O_2_ act as oxidative stress agent. However double mutant cells acted differently in response to genotoxic stress and microtubule inhibitor drug. The double mutant cell also showed the elevated Ty1 retro-transposition and multi nuclei phenotype indicating the genome instability phenotype. Further the mechanism of stress tolerance exhibited by *saf1Δrlm1Δ* cells needs further studies. However present study suggest a model (**Figure 11**) where WT cell shows sensitivity to the stress agents due to increased expression of RLM1 transcription factor which regulates expression of SAF1(positively) and AAH1 (negatively) genes during general stress. The ablation of both RLM1 and SAF1 together contributed to stress resistance due to loss of signaling. Based on the *in-silico* data analysis and experimental validation we suggest that RLM1 is novel transcription factor which regulates the expression of SAF1 and AAH1.

**Figure 11:**
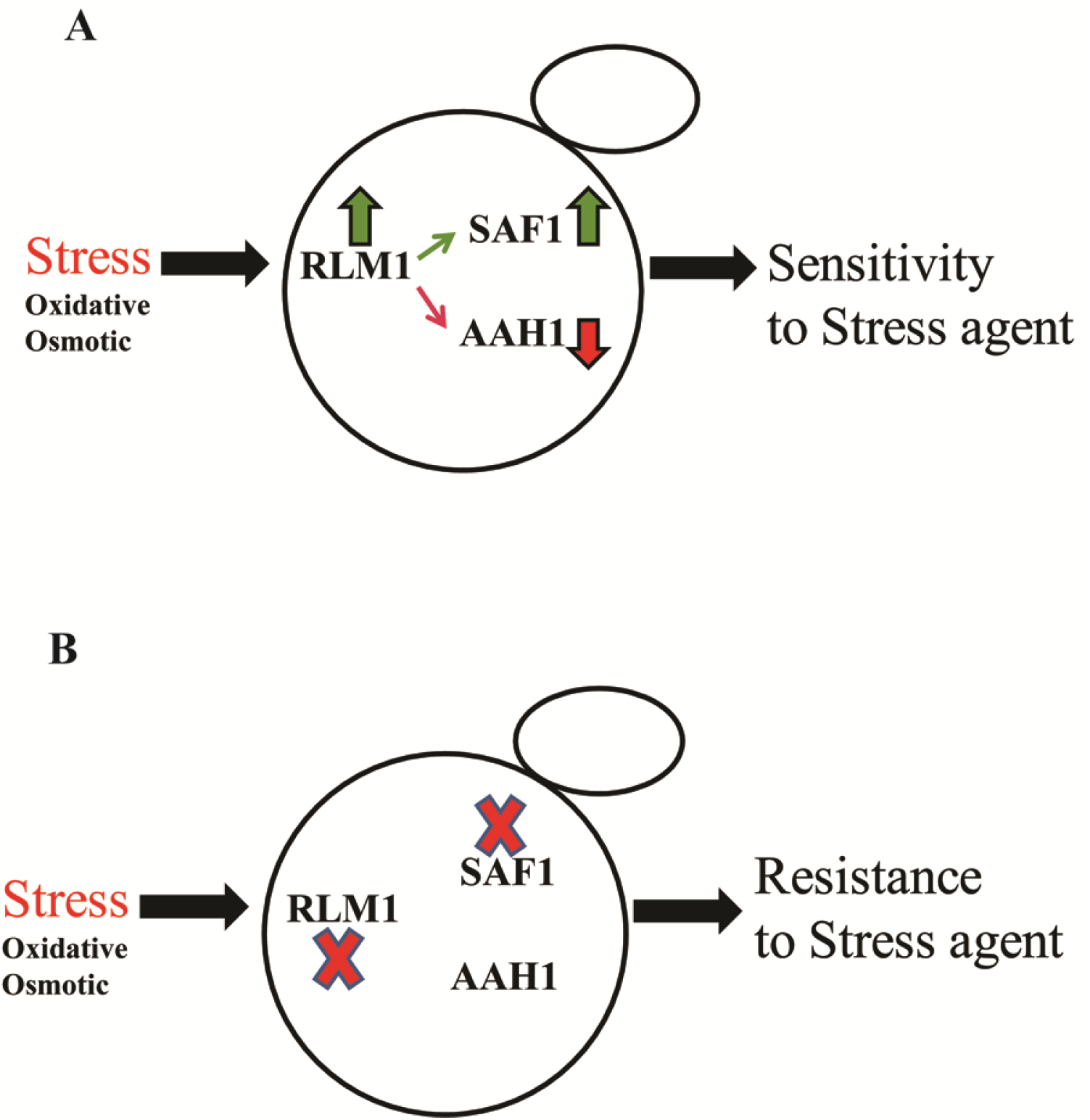
Model indicating the growth phenotype upon stress in WT and*saf1Δ rlm1Δ* cells. **A**. Stress mediated upregulation of RLM and SAF1 may lead to sensitive phenotype to stressor agent in WT cells. **B**. The ablation of both RLM1 and SAF1 genes together lead to resistance phenotype in presence of general stressor.

## Acknowledgment

We thank to Prof. M. Joan Curio, Dr. Deepak Sharma, IMTECH for strains and plasmids. We also thank Dr. Jitendra Thakur, NIPGR for lab support.

## Author’s contributions

NKB conceived and directed the study and wrote the paper with MS and VV. NKB and MS studied the database and analysed the *in-silico* database. MS performed the experiments and analysed data with NKB. VV provided the bioinformatics facility and analysed the data. All the authors reviewed the results and approved the final version of manuscript.

---

**Figure S1:**
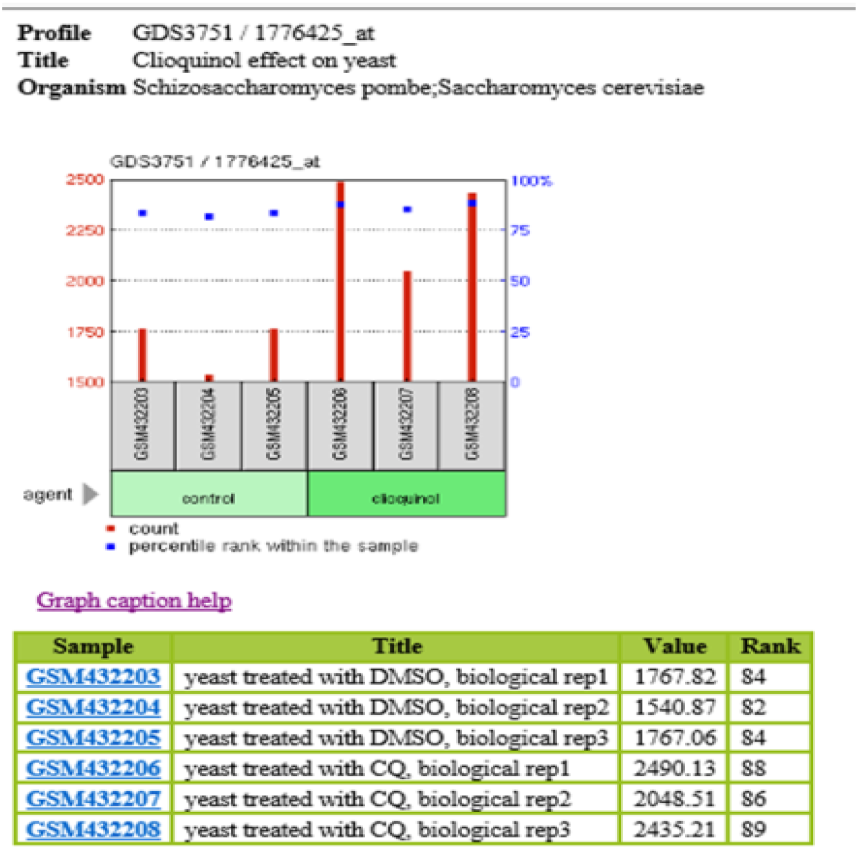
SAF1 overexpressed during stress in the WT cells. GEO profile of SAF1 in WT cells treated with Clioquinol, Pterostilbene, Gentamicin, Hypoxia condition, Genotoxic Stress, Desiccation, and Heat shock.

**Figure S2:**
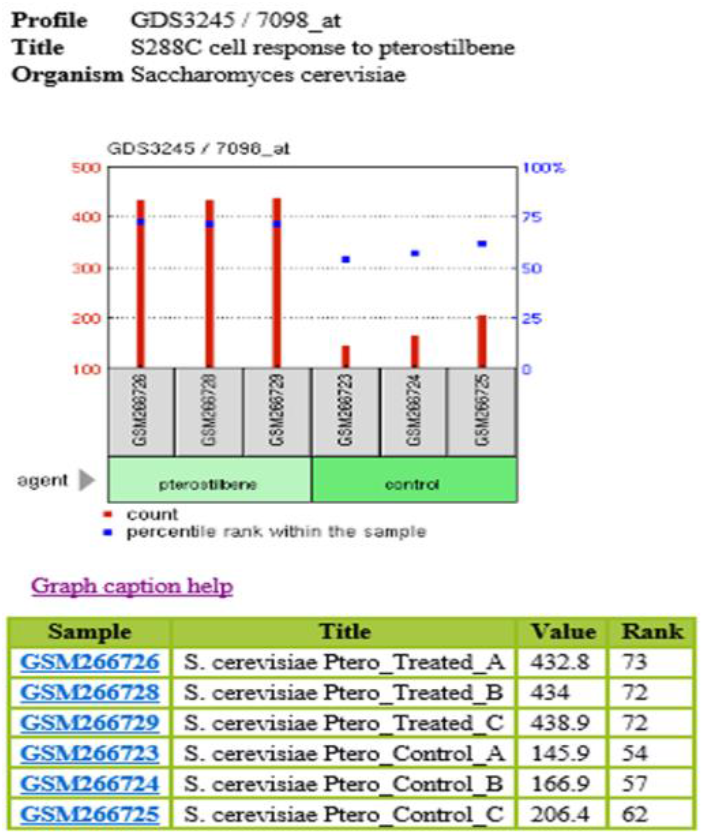
SAF1 overexpressed during stress in the WT cells. GEO profile of SAF1 in WT cells treated with Clioquinol, Pterostilbene, Gentamicin, Hypoxia condition, Genotoxic Stress, Desiccation, and Heat shock.

**Figure S3:**
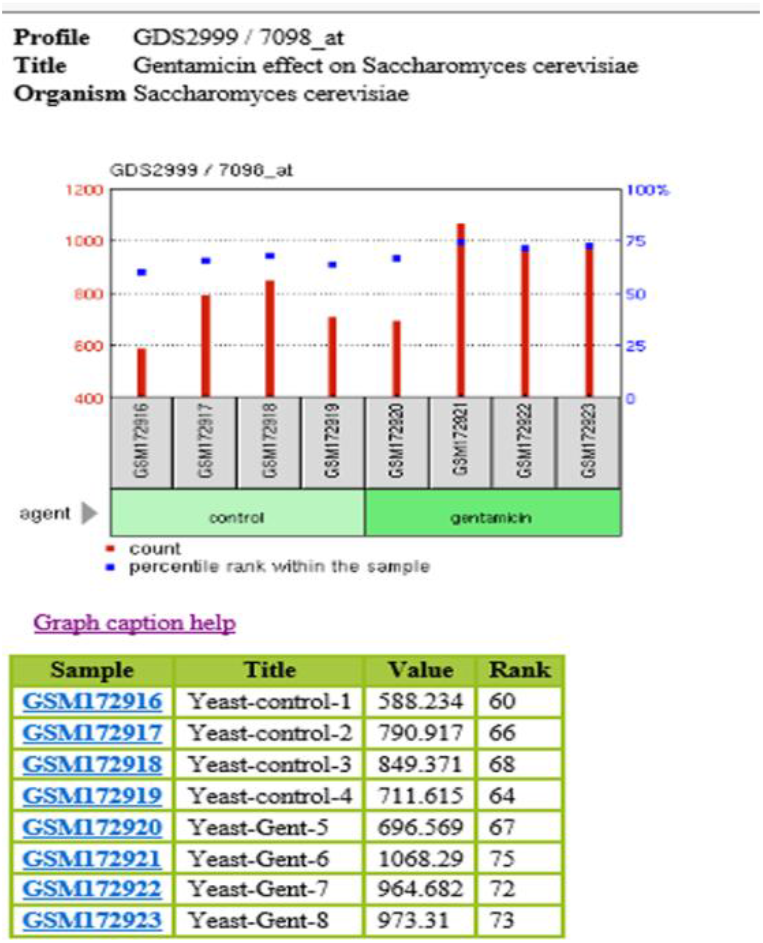
SAF1 overexpressed during stress in the WT cells. GEO profile of SAF1 in WT cells treated with Clioquinol, Pterostilbene, Gentamicin, Hypoxia condition, Genotoxic Stress, Desiccation, and Heat shock.

**Figure S4:**
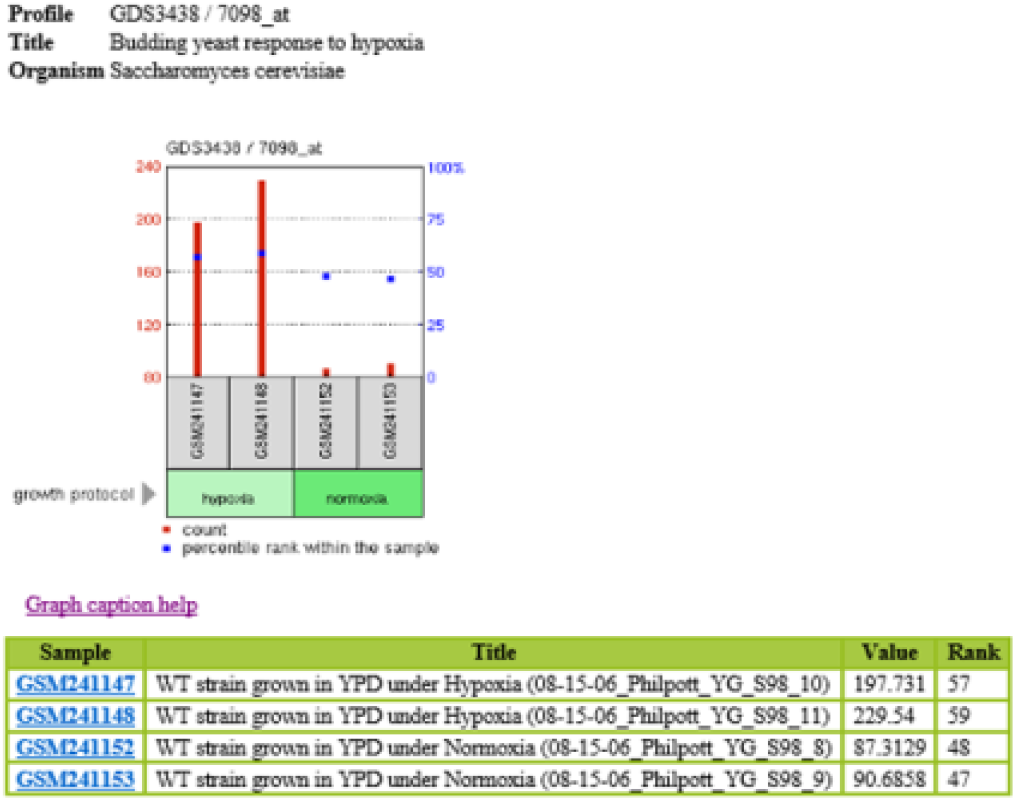
SAF1 overexpressed during stress in the WT cells. GEO profile of SAF1 in WT cells treated with Clioquinol, Pterostilbene, Gentamicin, Hypoxia condition, Genotoxic Stress, Desiccation, and Heat shock.

**Figure S5:**
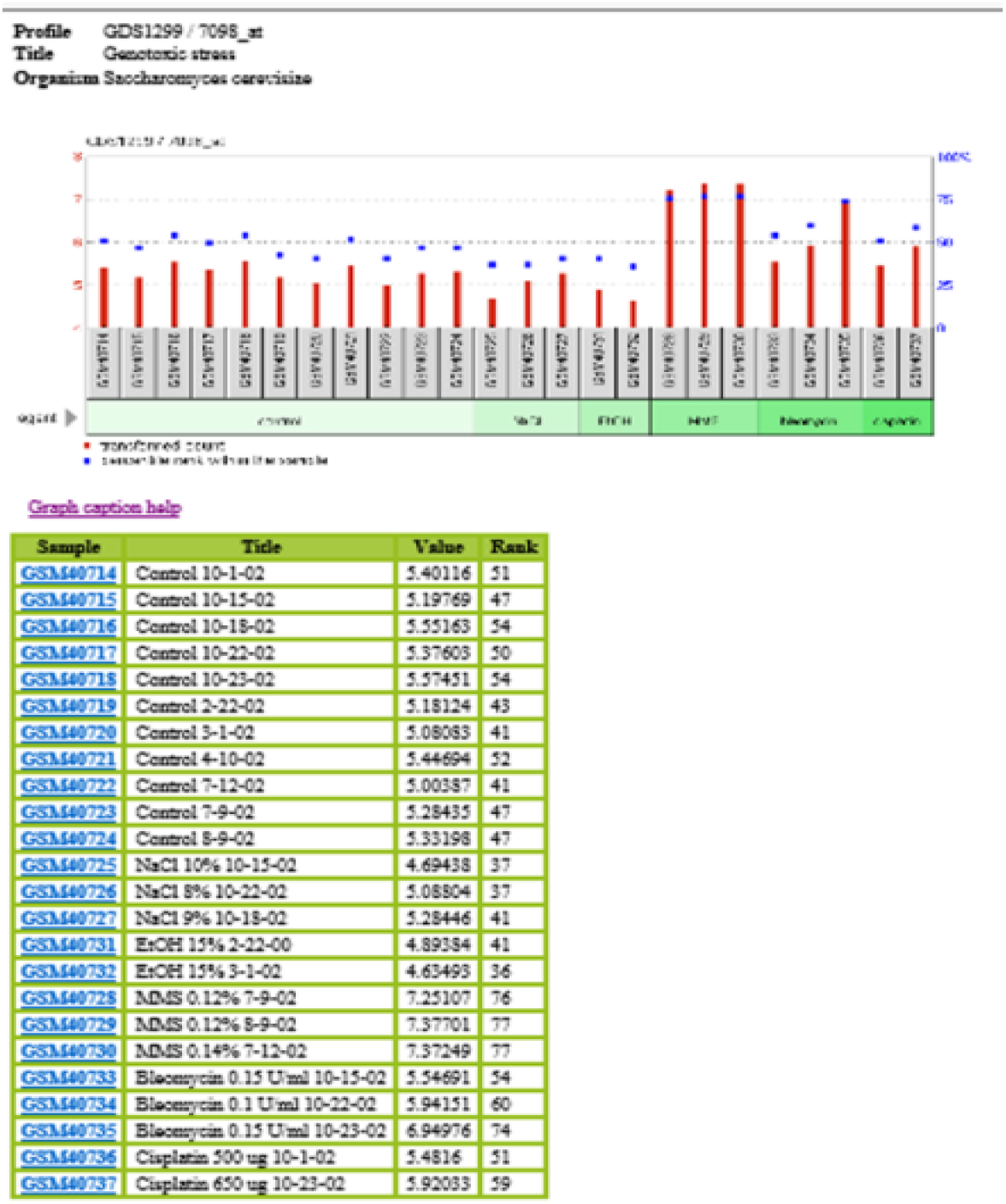
SAF1 overexpressed during stress in the WT cells. GEO profile of SAF1 in WT cells treated with Clioquinol, Pterostilbene, Gentamicin, Hypoxia condition, Genotoxic Stress, Desiccation, and Heat shock.

**Figure S6:**
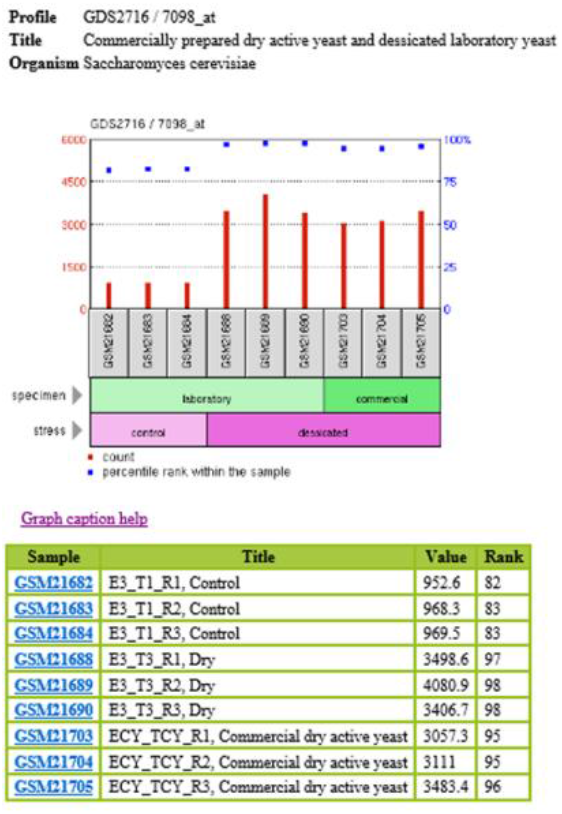
SAF1 overexpressed during stress in the WT cells. GEO profile of SAF1 in WT cells treated with Clioquinol, Pterostilbene, Gentamicin, Hypoxia condition, Genotoxic Stress, Desiccation, and Heat shock.

**Figure S7:**
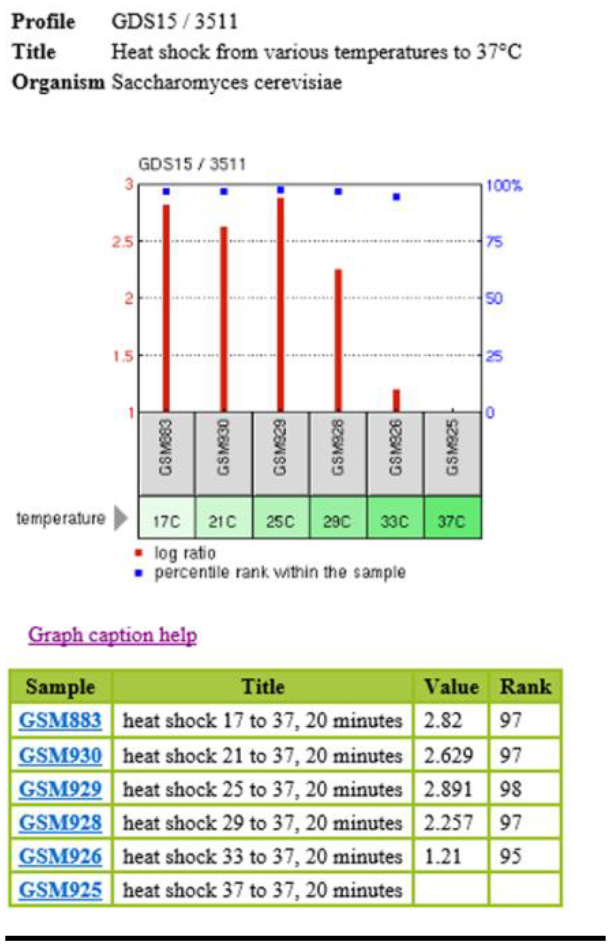
SAF1 overexpressed during stress in the WT cells. GEO profile of SAF1 in WT cells treated with Clioquinol, Pterostilbene, Gentamicin, Hypoxia condition, Genotoxic Stress, Desiccation, and Heat shock.

